# Improving allometric models to estimate the proboscis length of tropical bees

**DOI:** 10.1101/2025.02.21.639535

**Authors:** Kilian Frühholz, Kenneth Kuba, Maximilian Pitz, Julia Windl, Alexander Keller, Gunnar Brehm, Claus Rasmussen, Sara Diana Leonhardt, Ugo Mendes Diniz

## Abstract

The proboscis length of bees is a key morphological trait shaping communities, pollination networks, and likely their responses to habitat loss. Despite its importance, it is rarely considered in ecological studies because of logistic limitations in obtaining accurate measurements across many different species. In two previous studies, the proboscis length of temperate bee species was estimated based on body size and bee family. However, bee taxa partially occurring in the tropics might deviate from this allometric relationship due to different functional constraints. Thus, we tested if equations developed for temperate bees can accurately predict the proboscis length in Meliponini, Euglossini (both Apidae), and Augochlorini (Halictidae), three ubiquitous and highly important tribes of tropical bees. We measured the intertegular distance (as a proxy of body size measurement) and the proboscis length of 892 specimens of 105 tropical species. We used these measurements to evaluate the previous model and found that its estimations lacked accuracy when applied to tropical bees, particularly to Meliponini and Euglossini. We developed new allometric equations estimating the proboscis length based on the intertegular distance, using (sub-) genera as an additional predictive variable to refine the estimations. We tested our approach by creating a test model for Meliponini, trained with only 80 % of the data, and evaluated this model using the remaining 20 %, resulting in a high accuracy of estimates. Our results shed additional light on the nature of the proboscis length-body size allometric relationship in tropical bees and provide a tool for future studies on the functional ecology of bees and their interactions with plants.

## INTRODUCTION

Functional traits determine interactions between various trophic levels and within ecosystems (Schleuning et al. 2023). Traits can e.g. shape the distribution (Pollock et al. 2012), dispersal ability and niche occupation (Jiang et al. 2018), resource use (Gravel et al. 2016), or ability of species to adapt to a changing climate (Heilmeier 2019). To fully understand these mechanisms, an understanding of the underlying traits is crucial (McGill et al. 2006). A common example of species interactions, largely shaped by different functional traits, is pollination. Among the many variables that shape the interaction between pollinators and plants, the shape and length of their mouthparts play a particularly important role: proboscis length affects flower selection (Temeles et al. 2009; Basari et al. 2021; Inouye 1980; Haverkamp et al. 2016), foraging efficiency, pollination effectiveness (Haverkamp et al. 2016; Borrell 2007; Harder 1983; Peat et al. 2005), and extinction risk of pollinators and their host plant species (Stang et al. 2007). In a community-wide context, it regulates resource partitioning between pollinator species (Inouye 1978; Ranta 1984; Ranta and Lundberg 1980; Brown and Bowers 1985) and can be a driver of plant speciation (Borrell 2005; Rodríguez-Gironés and Santamaría 2007; Vajna et al. 2021). Thus, proboscis length is a key interaction trait that influences not only community assembly but also the structure of pollination networks by niche partitioning and determining patterns of specialization (Harmon-Threatt and Ackerly 2013; Stang et al. 2006, 2007; Stang et al. 2009).

For taxa such as hummingbirds or hawkmoths, whose long and often widely varying bills and proboscises indicate a clear functional specialization towards long-tubed flowers, the mouthpart length is routinely included in ecological studies that investigate the functional composition of these taxa (Torres-Vanegas et al. 2021), their interactions with plants (Guevara et al. 2023; Johnson et al. 2017) or the degree of specialization (Rodríguez-Flores et al. 2019; Nilsson and Rabakonandrianina 1988). Bees, however, despite being one of the most ubiquitous and ecologically dominant group of pollinators worldwide (Potts et al. 2010), are seldom explored in terms of the functional significance of their mouthparts. The few existing studies are mostly focused on large-bodied species like bumblebees (Goulson et al. 2008; Harder 1983; Stout et al. 2000). However, measuring the proboscis length of smaller bees usually requires dissection of the mouthparts, which can lead to destruction of the specimen, particularly in very small species, which then have to be identified beforehand (Cariveau et al. 2016). Thus, measuring the proboscis length of bees often is not feasible and multiple approaches have been taken to forego measuring proboscises in ecological studies, including adopting length categories, i.e., “long-tongued”, which include Apidae and Megachilidae, and “short-tongued” bees, including Halictidae, Andrenidae, Colletidae, Melittidae, and Stenotritidae (Michener 2007). This approach, however, lacks accuracy and brushes over taxon-specific differences (Ostwald et al. 2024). Thus, effort has been made to estimate proboscis length by allometric power functions including body size and bee family (Cariveau et al. 2016; Melin et al. 2019). Allometric functions can be used to describe the relationship between body size and metabolic rate, growth, or the size of specific body parts (Pélabon et al. 2014). Cariveau et al. (2016) and Melin et al. (2019) showed that the proboscis length of bee species increases with their body size in all families except the Australian Stenotritidae, which were not included in these studies. The proboscis length also differed between families, indicating that there is a phylogenetic component affecting the proboscis length of bees (Cariveau et al. 2016; Melin et al. 2019). However, these studies were carried out in temperate and subtropical regions, while data for tropical bees is lacking. Tropical bees may deviate from these allometric relationships due to functional constraints as a mechanism to avoid interspecific competition in these highly diverse ecosystems (Borrell 2005; Ostwald et al. 2024). Notably, morphological measurements to calculate functional diversity in tropical regions play an important role in the context of ongoing deforestation and its impact on global biodiversity (Wright and Muller-Landau 2006; van der Sluijs 2020; Alroy 2017; Ostwald et al. 2024). This is especially true for monitoring the effects of forest restoration as patterns of resistance and recovery of insect communities are often tied to dispersal and interaction traits (D’Astous et al. 2013; Montoya-Pfeiffer et al. 2018; Audino et al. 2014; Montoya-Pfeiffer et al. 2020; Lichtenberg et al. 2017). Functionally diverse pollinator communities ensure pollination services within the process of tropical forest recovery and conservation. Hereby, bees play an essential role as they are responsible for the pollination of the majority of tropical plants (Michener 2007; Ollerton et al. 2011). It is therefore paramount to streamline and standardize the estimation of proboscis length.

In the Neotropics, three of the most abundantly encountered bee tribes are Meliponini (Apidae), Euglossini (Apidae), and Augochlorini (Halictidae) (Michener 2007). Systematic studies on the morphological traits of these bee tribes are scarce. Although the proboscis length is used as a trait to identify Euglossini males (Bembé 2007) and is comparatively easy to measure because of its length, it is rarely considered in ecological studies (e.g., Brito et al. 2018; Guevara et al. 2024). There are a few studies which have assessed the morphometrics of single Meliponini species (Basari et al. 2021; Kiatoko et al. 2023), and some have further placed the proboscis length into an ecological context to uncover floral preferences (Laha et al. 2020) or to explore the phenology of plant-pollinator interactions (Ribeiro et al. 2024).

The importance of the proboscis length as a community-shaping morphological trait (Harmon-Threatt and Ackerly 2013) makes it necessary to get the most accurate values, which is achieved by direct specimen measurements. However, in large biodiversity assessments with a multitude of species, direct measurements might be constrained by time and resources and may need to be replaced by estimates. Expanding the existing models to tropical species might therefore not only allow such estimations, but also add to our understanding of general allometry in bees, which contributes to recognizing species that differ from expected patterns and explaining certain life history traits (Pélabon et al. 2014).

We thus aimed to assess the allometric relationship between bee size and proboscis length in these key tropical bee tribes (Meliponini, Euglossini, and Augochlorini) by (i) testing the applicability of pre-existing models on a large database of bees caught in a diverse lowland rainforest ecosystem in Ecuador and (ii) providing updated model versions for tropical bees that also account for bee tribe and genus. We hypothesized that the proboscis length of the two tropical bee tribes which were not included in the dataset of Cariveau et al. (2016), i.e. Meliponini and Euglossini, could not be accurately predicted by the existing model, while it would provide accurate estimates for the proboscis length of Augochlorini. Based on our findings, we furthermore composed an R package expanding the scope of the previous model to easily estimate the proboscis length of tropical bees using measurements of intertegular distance and taxonomic information.

## METHODS

### Data collection

Specimen collection was carried out between March and December 2023 in the Reserva Río Canandé (0°31’33.4”N, 79°12’46.0”W) and Reserva Tesoro Escondido (0°32’30.9”N, 79°08’41.9”W), northwestern Ecuador, Chocó-Darien ecoregion. Specimens were collected via fragrance traps aimed at Euglossini males (Ferreira et al. 2013), vane traps with blue and yellow vanes (Prendergast et al. 2020; Renteria and Brehm 2025), and active netting. All bees were killed with chloroform fumes. Methods are described in more detail in Diniz et al. (2025) and Escobar et al. (2024).

After collection, all specimens were identified to the lowest possible taxonomic level using specialized keys (Engel et al. 2023; Engel 2000; Bonilla-Gómez and Nates-Parra 1992). Intertegular distance (IT), defined as the distance between the tegulae, was used as a body size proxy (Cariveau et al. 2016; Stemet et al. 2024; Kendall et al. 2019). The proboscis length (PB) was defined as the length of the prementum plus the length of the glossa or the distance from the base of the mentum to the distal point of the labellum (Cariveau et al. 2016) (Supplementary Information, Fig. S1). Measurements from the smaller sized Meliponini and Augochlorini were done in photos taken with a Wild Heerbrug M7S microscope attached to a Leica MC120 HD camera using ImageJ (Schneider et al. 2012). For Meliponini and Augochlorini, the proboscis was dissected and mounted on slides for photographing. Measurements of the comparatively large Euglossini bees were taken with a caliper on fully stretched and straight proboscises. All measurements were taken on specimens which had been stored in 70 %-Ethanol at −20 °C. All measured Meliponini were female workers, Augochlorini were females, and Euglossini were males. Cariveau et al. (2016) found no differences in PB-allometry between sexes.

### Data analysis

All analyses were performed in R 4.4.1 (R Core Team 2024). First, to test the quality of pre-existing allometric model, PB was estimated via IT using the *BeeIT* package (Cariveau et al. 2016). The estimated values were compared to the observed values and model fit was assessed via linear regression using R² and root mean squared error (RSME) as comparative metrics, with a high R² and a low RSME indicating a good fit.

Then, four new linear models were created separately for each bee group, including IT and (sub-) genus as predictive variables independently and as interacting parameters. For easier application and to include lower taxonomic levels than family, we adapted the allometric equation as follows:

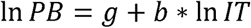

where g = a coefficient specific to the genus, subgenus or tribe and b = an allometric scaling coefficient, representing the slope of the model. The models’ relative fit was evaluated using the Akaike information criterion (AIC). Unlike Cariveau et al. (2016) and Melin et al. (2019) who calculated species trait means, we used all individual measurements to account for interspecific variation, to increase the overall amount of training data, especially as some species were represented by only one specimen and to account for groups with difficult separation of biological species, e.g. Augochlorini.

To evaluate the accuracy of our approach, we created a second model by training it with only 80 % of the measured Meliponini data to estimate PB. We then used the other 20 % to estimate model fit and calculated the difference between the predicted and the measured values and the 90 %-quantile of this difference as a metric for the accuracy of our estimates.

## RESULTS

In total, 892 specimens from 105 species were measured (1 to 36 individuals per species): 504 Euglossini from 58 species, 343 Meliponini from 36 species, and 44 Augochlorini from 11 species (Supplementary Information, Table S2). From 16 species only one individual was available for measurement.

### Evaluation of the pre-existing model

When fit to our data, the model from Cariveau et al. (2016) did not provide accurate estimates of proboscis length for the three investigated bee tribes. The proboscis length of Meliponini was overestimated by 0.71 mm on average (sd = 0.49 mm) and residuals ranged from −0.71 to 1.96 mm (max. deviation > 123 % of IT), which resulted in a low R² and a high RMSE (Table 1). The proboscis length of Euglossini was greatly underestimated with a mean residual of −7.08 mm (sd = 6.24 mm) and residuals ranging from −25.83 to 3.48 mm (max. deviation > 759 % of IT), which resulted in a negative R² and a high RMSE (Table 1). Of the three tribes, the model most accurately predicted the proboscis length of Augochlorini with a mean residual of −0.07 mm (sd = 0.35 mm) and a range of residuals from −1.04 to 0.58 mm (max. deviation > 85 % of IT), which resulted in a low RMSE (Table 1).

**Table 1.**
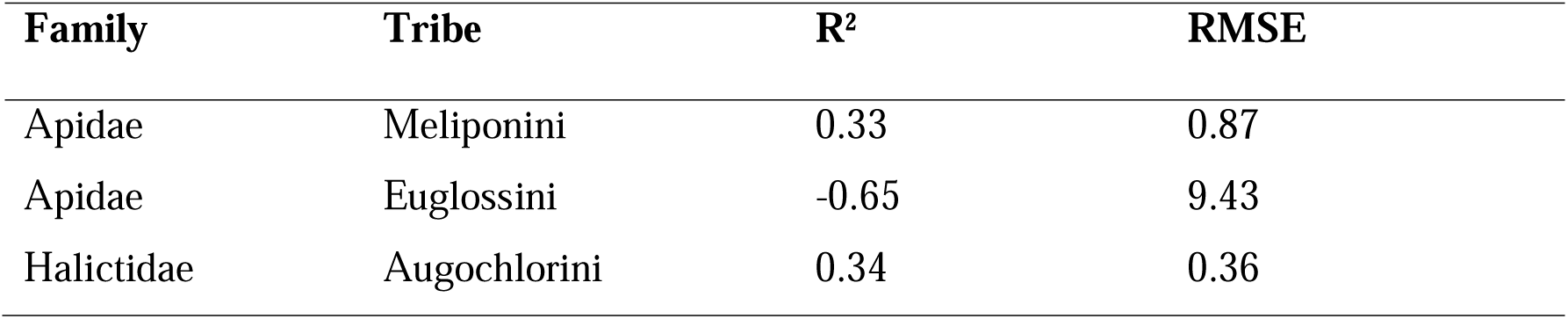
Goodness of fit measures for the model from Cariveau et al. (2016) against the measured.

### Improved allometric models

All new linear models showed a better fit in comparison to the pre-existing model of Cariveau et al. (2016). The most parsimonious models for Augochlorini, Meliponini and all Euglossini included IT and genus as independent predictors, while an interaction between the two parameters showed a similar or better fit to the data (Table 2). For Euglossini (*Euglossa* excluded) and *Euglossa*, the models with the lowest AIC additionally included the interaction between IT and (sub-) genus. Including (sub-) genus as a predictive variable notably lowered model AIC in all groups, indicating that the mean proboscis length differs between (sub-) genera. For Meliponini and Euglossini, the models with only (sub-) genus had a notably lower AIC than the models including only IT (Table 2).

**Table 2.** Goodness of fit measures of the new linear models for every bee tribe and additionally for Euglossini excluding *Euglossa* and a model for *Euglossa* including subgenera. The best model with the lowest AIC is marked in bold.

| Tribe | Model | R<sup>2</sup> | AIC |
| --- | --- | --- | --- |
| Meliponini | IT | 0.71 | 48.39 |
|  | genus | 0.82 | -96.26 |
|  | IT + genus | **0.83** | **-114.02** |
|  | IT * genus | 0.85 | -112.22 |
| Euglossini | IT | 0.14 | 620.45 |
|  | Genus | 0.20 | 589.81 |
|  | IT + genus | **0.22** | **577.56** |
|  | IT * genus | 0.23 | 578.30 |
| Euglossini (without Euglossa) | IT | 0.10 | 105.56 |
|  | Genus | 0.65 | -15.03 |
|  | IT + genus | 0.65 | -12.90 |
|  | IT * genus | **0.68** | **-19.45** |
| Euglossini (only Euglossa) | IT | 0.04 | 504.94 |
|  | Subgenus | 0.72 | 47.51 |
|  | IT + subgenus | 0.72 | 49.27 |
|  | IT * subgenus | **0.76** | **-8.40** |
| Augochlorini | IT | 0.51 | -42.71 |
|  | Genus | 0.63 | -52.60 |
|  | IT + genus | **0.71** | **-60.73** |
|  | IT * genus | 0.74 | -60.39 |

Figure 1A shows the tribe-specific relationship between IT and PB without considering genus, while Figure 1B shows the relationship as indicated by the lowest AIC. The model for all Euglossini showed the worst fit, however, separating *Euglossa* and the other genera and including the subgenus in the Euglossa model increased the model fit greatly. Means for IT and PB for all species are provided in the supplementary material (S 2). The power function was parameterized to predict PB of Meliponini, Euglossini, and Augochlorini using the estimates of the models with the best fit (Table 3).

**Figure 1.**
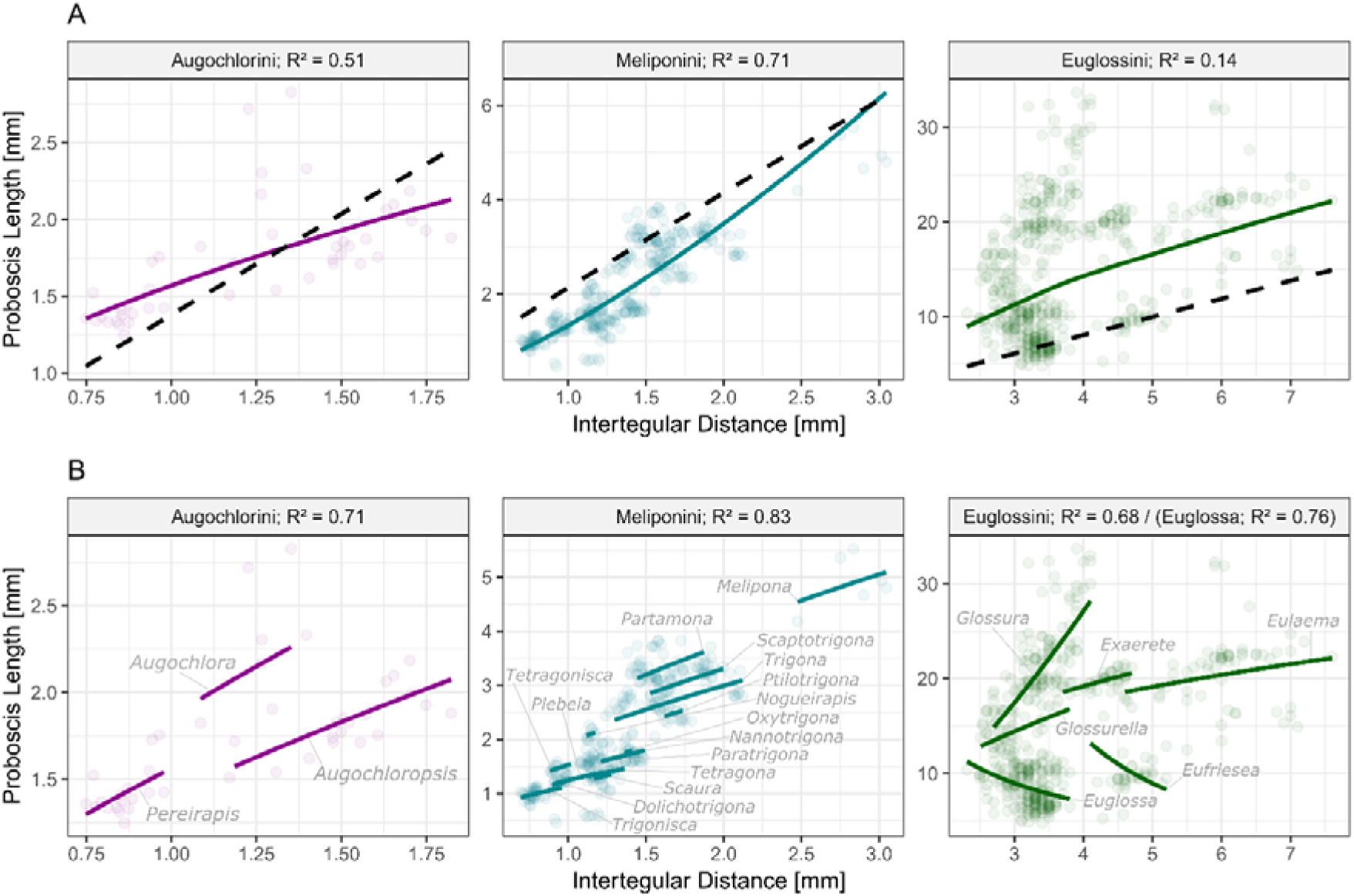
Relationship between intertegular distance and proboscis length for the three tribes (A) and separated between (sub-) genera (B) with R2-values provided for our models. Each point represents one measured specimen. The dashed lines in A represent the estimates by Cariveau et al. (2016). The full lines represent the estimates by the new models. In B the names of the genera are specified.

**Figure 2.**
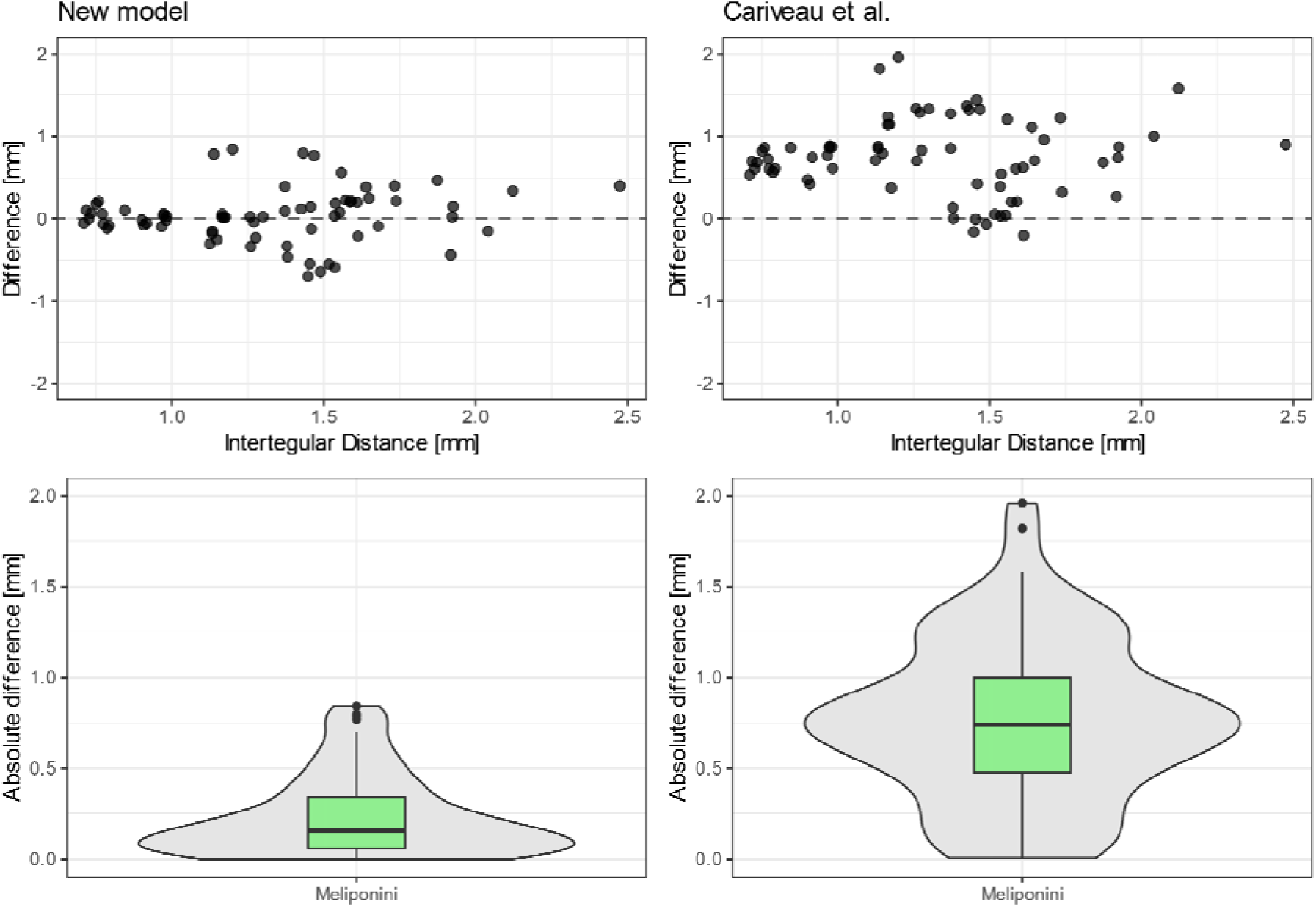
Error distribution of the new model (left) and the model of Cariveau et al. (2016) (right) when applied to the evaluation data from Meliponini.

**Table 3.**
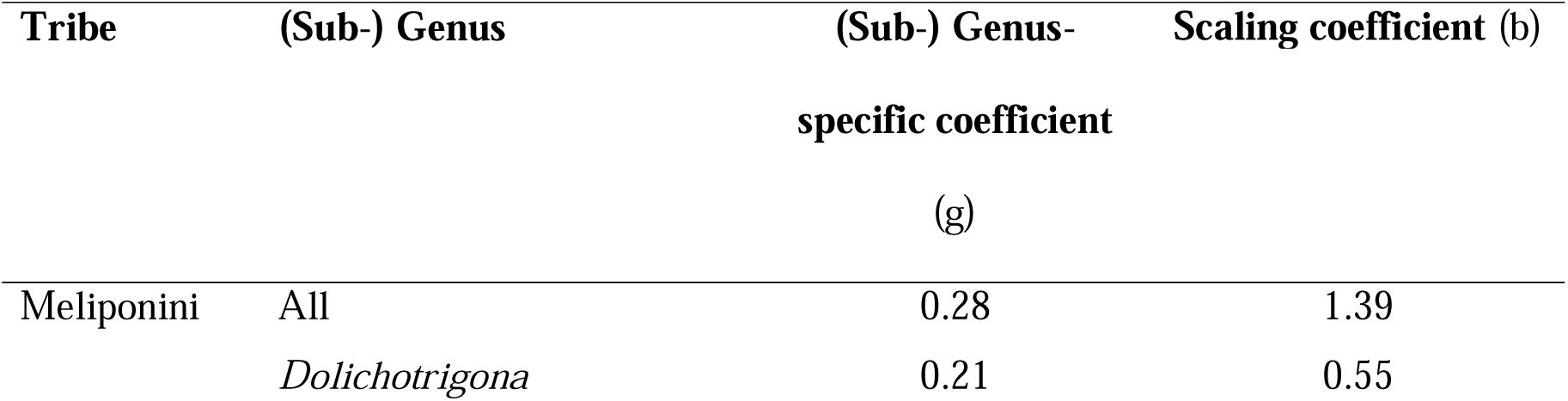
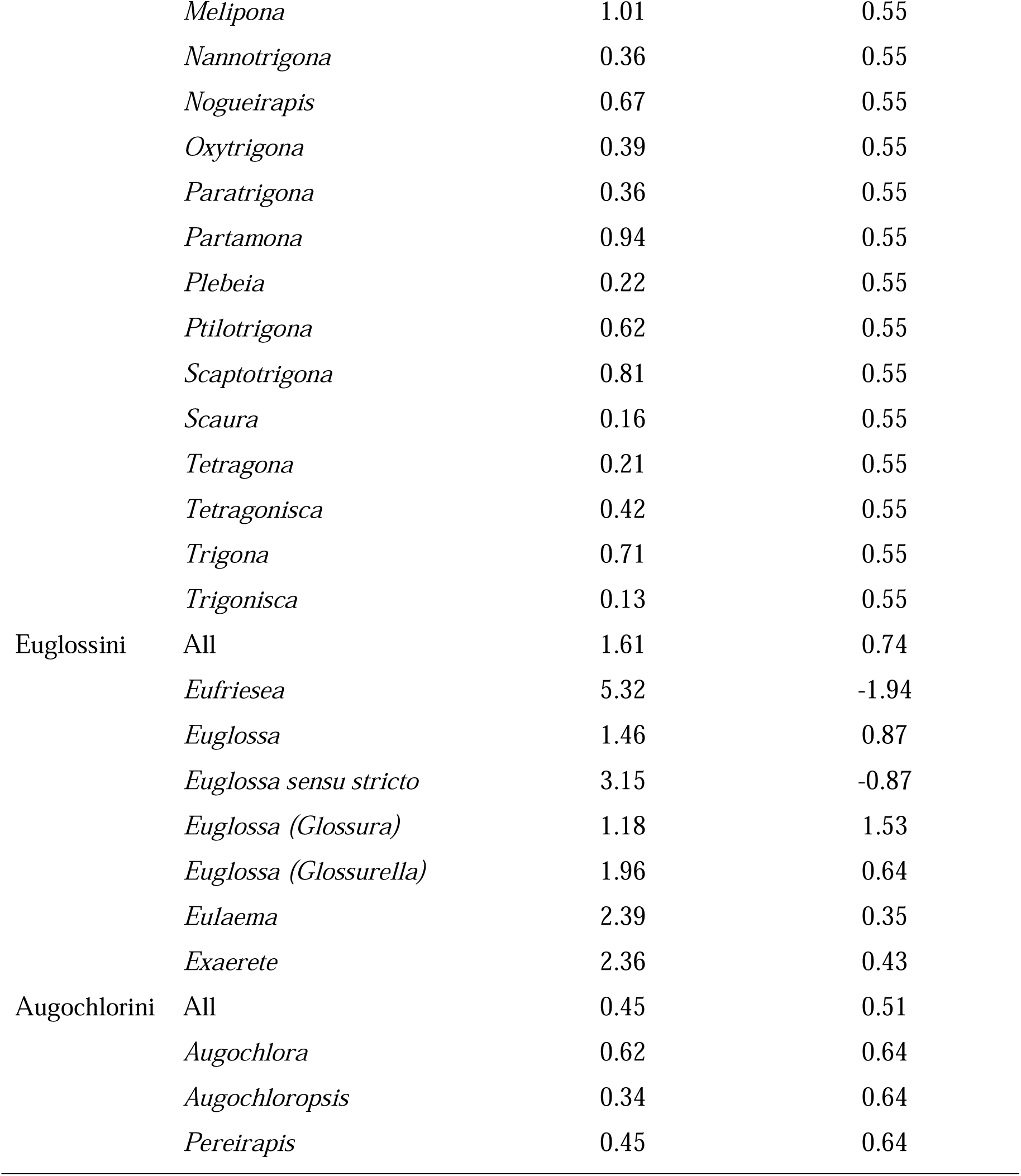
Coefficients of the allometric equations for the three bee tribes, parameterized by intercept (g) and slope (b) of the best linear model for each tribe and genus. Genus “All” specifies the model coefficients for the tribe-based model (PB ∼ IT) for genera not considered in this study. For *Eufriesea*, *Eulaema,* and *Exarete* the model excluding *Euglossa* was used. By inserting g and b in the formula, the proboscis length can be calculated.

| Tribe | (Sub-) Genus | (Sub-) Genus-specific coefficient (g) | Scaling coefficient (b) |
| --- | --- | --- | --- |
| Meliponini | All | 0.28 | 1.39 |
|  | *Dolichotrigona* | 0.21 | 0.55 |
|  | *Melipona* | 1.01 | 0.55 |
|  | *Nannotrigona* | 0.36 | 0.55 |
|  | *Nogueirapis* | 0.67 | 0.55 |
|  | *Oxytrigona* | 0.39 | 0.55 |
|  | *Paratrigona* | 0.36 | 0.55 |
|  | *Partamona* | 0.94 | 0.55 |
|  | *Plebeia* | 0.22 | 0.55 |
|  | *Ptilotrigona* | 0.62 | 0.55 |
|  | *Scaptotrigona* | 0.81 | 0.55 |
|  | *Scaura* | 0.16 | 0.55 |
|  | *Tetragona* | 0.21 | 0.55 |
|  | *Tetragonisca* | 0.42 | 0.55 |
|  | *Trigona* | 0.71 | 0.55 |
|  | *Trigonisca* | 0.13 | 0.55 |
| Euglossini | All | 1.61 | 0.74 |
|  | *Eufriesea* | 5.32 | -1.94 |
|  | *Euglossa* | 1.46 | 0.87 |
|  | *Euglossa sensu stricto* | 3.15 | -0.87 |
|  | *Euglossa (Glossura)* | 1.18 | 1.53 |
|  | *Euglossa (Glossurella)* | 1.96 | 0.64 |
|  | *Eulaema* | 2.39 | 0.35 |
|  | *Exaerete* | 2.36 | 0.43 |
| Augochlorini | All | 0.45 | 0.51 |
|  | *Augochlora* | 0.62 | 0.64 |
|  | *Augochloropsis* | 0.34 | 0.64 |
|  | *Pereirapis* | 0.45 | 0.64 |

Our test model aimed at predicting Meliponini PB was trained with 274 random measurements, while the other 69 data points were used as the evaluation data set. With our new model, absolute differences between the measured and estimated data points were always below 1 mm and errors were equally distributed around 0 mm (mean = 0.03 mm, sd = 0.32 mm). In contrast, differences between measured and estimated value using the Cariveau et al. (2016) model often exceeded 1 mm and tended to overestimate PB of Meliponini (mean = 0.75 mm, sd = 0.46 mm). For our new model, 90% of the absolute errors were below 0.57 mm, while the 90%-quantile of the absolute errors for the model of Cariveau et al. (2016) was at 1.33 mm. The new model accurately predicted PB particularly of small Meliponini, while errors were larger for medium-sized individuals.

For easier use, we implemented the allometric equations in the R package *tropTongue*, which can be downloaded from https://github.com/kilian-fru/tropTongue. The parameters used are specified in Table 3. For other tribes, the package uses the family-specific function from the package *BeeIT* (Cariveau et al. 2016).

## DISCUSSION

In this study, we showed that a pre-existing allometric model created for bees in temperate environments (Cariveau et al. 2016) could not accurately predict the proboscis length of several tropical bee tribes. New allometric models were created including the intertegular distance as body size proxy and the (sub-) genus. This significantly improved the accuracy of the models when compared to the Cariveau et al. (2016) model. Model outcomes confirmed an allometric relationship between body size, phylogeny and mouthpart length for the tropical bee tribes studied, which has also been found in other nectar-feeding animals, e.g. tropical butterflies (Kunte 2007), tropical Sphingidae (Agosta and Janzen 2005), subtropical and temperate wild bees (Menegus 2018; Cariveau et al. 2016; Melin et al. 2019), birds (Rombaut et al. 2022) and non-nectar-feeding animals, e.g. weevils (Fleurot et al. 2022) and certain tropical butterfly clades (Kunte 2007).

We showed that the allometric relationship between body size and proboscis length varies between temperate and tropical bees, at least in Apidae (e.g., Meliponini, Euglossini). Having an overproportionately long proboscis, like in the case of many Euglossini compared to temperate Apidae, is usually considered a competitional advantage as plants with short- and long-tubed flowers both can be used for foraging (Borrell 2005). However, long-tongued insects were found to exhibit longer flower-handling times and higher energy use while foraging (Kunte 2007; Harder 1983; Borrell 2007). They thus likely target longer-tubed flowers which offer a higher amount of nectar while excluding insects with shorter proboscises (Dressler 1982; Johnson et al. 2017). On the other hand, very small bees, like Meliponini, might be able to compensate for their shorter-than-expected proboscis by their small body size, making them able to crawl into narrow-tubed flowers to forage (Engel et al. 2023; Michener 2007).

Extreme values for proboscis length like those described but also for other functional bee traits are more likely to appear in the tropics: Tropical ecosystems are the most biodiverse on Earth, harboring not only a high plant species diversity (Myers et al. 2000) but also an even higher functional diverse flora than their species pool would suggest (Swenson et al. 2012). The functional diversity of plants increases towards the equator (Lamanna et al. 2014), due to for example a high number of epiphytes and lianas (Spicer et al. 2020), which might affect the functional diversity of bees. Proboscis length is an important interaction trait linked to the morphological matching between plants and pollinators (Goulson et al. 2008). Because of the high (functional) plant diversity, the trait space occupied by tropical bee communities is likely also larger than the space occupied by temperate bee communities, resulting in a wider range of proboscis lengths in tropical bees.

Studying functional traits in these highly diverse ecosystems is challenging. The approach to estimate difficult-to-measure morphological traits, like the proboscis length, by allometric equations can greatly simplify data collection (Cariveau et al. 2016; Ostwald et al. 2024). However, as we showed in our study, estimated values should be handled carefully. Cariveau et al. (2016) pointed out, that adopting length categories (“long-tongued” / “short-tongued”) to overcome measurements of proboscis length does not provide sufficient accuracy and thus introduced an allometric equation to estimate the proboscis length using body size and bee family, parameterized with measurements of temperate bees. Our results suggest that combining body size measurements at lower taxonomic levels can further improve estimates of bee proboscis length, at least in the tropics (Ostwald et al. 2024). Especially in morphologically highly diverse families such as Apidae (Engel et al. 2021), we suggest to include the tribe or the genus to increase accuracy, wherever it is feasible. Including the subgenus in the equation further improved estimates and model fit in *Euglossa*, whose subgenera are partly separated by proboscis length (Bonilla-Gómez and Nates-Parra 1992; Bembé 2007) suggesting a strong phylogenetical component shaping the differentiation of proboscis length in this genus. *Euglossa* represents the largest genus of Euglossini with 139 often co-existing species (Engel and Rasmussen 2019). Differentiation in proboscis length might thus also be a strategy to overcome competition between species within *Euglossa*.

Further model adaptations might focus on enhancing data availability and thus quality for tropical bees. Additional data would be especially useful in evaluating such allometric models. The evaluation of our approach with the test model showed an increased accuracy of our model for Meliponini, however, using independent data is crucial to confirm our and future findings. Further improving data quality through e.g. incorporating data from other studies will require a standard protocol to measure proboscis length of bees (Keller et al. 2023). For example, we would have liked to add data from Ribeiro et al. (2024), but they used another measuring protocol preventing comparison of measurements. To improve data availability we used individual measurements instead of species means, in contrast to previous models (Cariveau et al. 2016). This approach might bias our allometric equations towards more abundant species, which were measured most frequently. It does however increase the overall quality of the model, especially for groups like Augochlorini, where species identification is difficult and not many individuals were available, rendering means even less accurate. Studies on tropical Augochlorini are scarce and taxonomic keys are missing, even though they might be important indicators of forest loss as they appear to depend on unforested habitats for nesting (Brosi et al. 2007). Additionally, individual-based measurements were shown to produce the same results as species means while decreasing measuring effort and enhancing data availability (Beck et al. 2024).

The most accurate method to obtain values of proboscis length is to measure them manually. However, in large biodiversity assessments, for large species pools, very small or rare species or species with unclear taxonomic status, like often found in the tropics, this is not feasible. Our model enables researchers to use body size measurements to additionally infer proboscis length, when it could not be manually obtained. Body size is measured comparatively often in ecological studies on bees (Osorio-Canadas et al. 2022; Lichtenberg et al. 2017; Montoya-Pfeiffer et al. 2020) but the proboscis length is rarely considered.

Our results shed additional light on the nature of the proboscis length-body size allometric relationship in tropical bees and may serve as an additional tool for future ecological studies that want to include proboscis length to assess bee (functional) diversity, morphology and allometry in the tropics. The proboscis length of bees is related to various aspects of bee ecology such as flower selection (Basari et al. 2021) and competition (Ranta and Lundberg 1980), determines patterns of distribution and habitat preferences (Harmon-Threatt and Ackerly 2013) and might be linked to pesticide uptake by bees (Kopit and Pitts-Singer 2018; Borrell 2007). Therefore, it directly affects the conservation of tropical bee communities which are mostly endangered by habitat loss in particular deforestation and pesticides (Toledo-Hernández et al. 2022). Furthermore, proboscis length has a strong effect on the pollination services of bees (Chase et al. 2023), making it an important trait to consider when assessing tropical forest restoration and conservation, which is largely influenced by bee pollination (Ollerton et al. 2011). We, hereby, emphasize the importance of including the proboscis length in much needed research on tropical bee functional ecology, which is crucial in understanding the effects and underlying patterns of tropical deforestation, forest restoration, biodiversity loss and climate change.

## ACKNOWLEDGEMENTS

This work was funded by the Deutsche Forschungsgemeinschaft (DFG) funded Research Unit REASSEMBLY (FOR 5207; sub-projects LE2750/12-1 and KE1742/13-1). We thank the Ministry of Environment of Ecuador for granting research and collection permits through Contrato Marco MAE-DNB-CM-2021-0187, Sebastián Escobar for handling exportation permits, Martin Schaefer (Fundación Jocotoco) and Citlalli Morelos-Juarez (Fundación Tesoro Escondido) for allowing us to work in their reserves, and the staff of both reserves: Katrin Krauth, Julio Carbajal, Jender Vélez, Bryan Tamayo, Lady Condoy, Leonardo de la Cruz, Jefferson Tacuri, Yadira Giler and Adriana Argoti.

## AUTHOR CONTRIBUTIONS

KF, KK, SDL and UMD conceptualized the research. SDL, AK and GB acquired and managed the funding. UMD, SDL and JW performed fieldwork and data collection. KF, KK, MP, JW and UMD did sample processing and data curation. CR, KF, MP, JW and UMD identified insects. KF performed data analysis and wrote the manuscript draft. All authors contributed critically to the last manuscript draft.

## CONFLICT OF INTERST

The authors declare no conflicts of interest.

## DATA AVAILABILITY

Data and R-code are available under https://github.com/kilian-fru/tropTongue-analysis.

