## Supplementary figures and images for "Improving allometric models to estimate the proboscis length of tropical bees"

### Figure S1

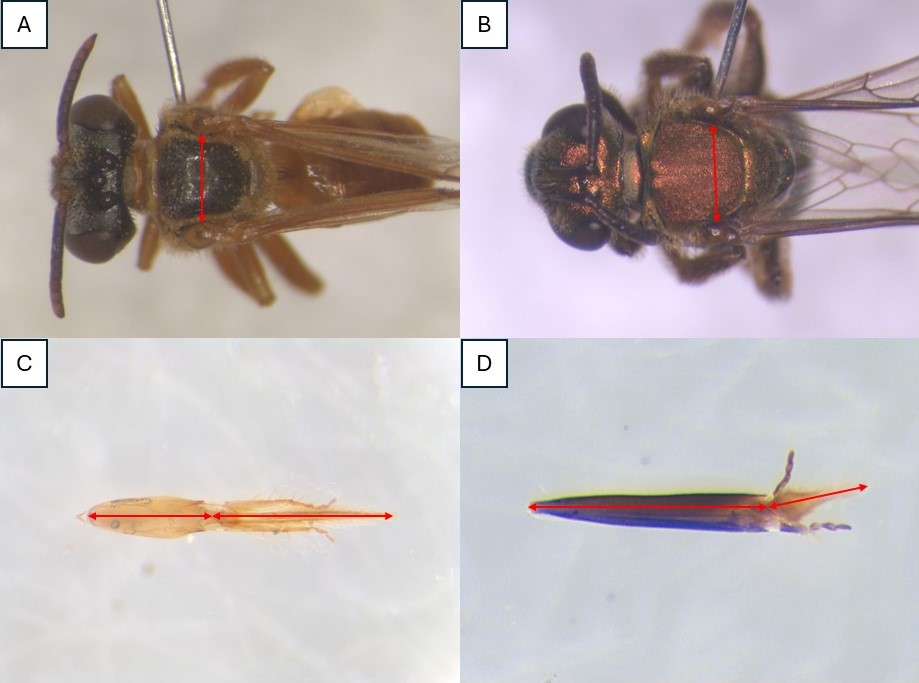
